# Fast and sensitive protein sequence homology searches using hierarchical cluster BLAST

**DOI:** 10.1101/426098

**Authors:** Daniel J. Nasko, K. Eric Wommack, Barbra D. Ferrell, Shawn W. Polson

## Abstract

The throughput of DNA sequencing continues to increase, allowing researchers to analyze genomes of interest at greater depths. An unintended consequence of this data deluge is the increased cost of analyzing these datasets. As a result, genome and metagenome annotation pipelines are left with a few options: (*i*) search against smaller reference databases, (*ii*) use faster, but less sensitive, algorithms to assess sequence similarities, or (*iii*) invest in computing hardware specifically designed to improve BLAST searches such as GPGPU systems and/or large CPU-rich clusters.

We present a pipeline that improves the speed of amino acid sequence homology searches with a minimal decrease in sensitivity and specificity by searching against hierarchical clusters. Briefly, the pipeline requires two homology searches: the first search is against a clustered version of the database and the second is against sequences belonging to clusters with a hit from the first search. We tested this method using two assembled viral metagenomes and three databases (Swiss-Prot, Metagenomes Online, and UniRef100). Hierarchical cluster homology searching proved to be 12-times faster than BLASTp and produced alignments that were nearly identical to BLASTp (precision=0.99; recall=0.97). This approach is ideal when searching large collections of sequences against large databases.

## Background

Advancements in DNA sequencing continue to have a profound impact in biology and has revitalize the field of genomics. The cost of DNA sequencing has fallen dramatically over the last 10 years and the throughput has increased at a rate that has greatly exceeded Moore’s law [1]; Gordon Moore’s axiom which has accurately predicted the rate of advancement for computational hardware over the last forty years. Not only are the size of sequencing datasets increasing (e.g. a dual S4 flow cell run on the NovaSeq 6000 System generates up to six tera base pairs per run), but large reference databases (e.g. RefSeq, UniRef) are doubling in size every two years. At its current rate, by the year 2024, UniRef100 may contain over one billion peptide sequences that total nearly half a trillion amino acids.

The CPU requirements of homology searches against large reference databases are the primary computational constraint in genome and metagenome annotation pipelines. The Smith-Waterman algorithm [2] was among the first algorithms capable of searching for homology between two sequences. Smith-Waterman is guaranteed to produce the optimal local alignment of any two sequences, but it is far too slow, even when searching a small set of experimental sequences against a small- to medium-sized set of known reference sequences. Heuristic algorithms such as FASTA [3] and BLAST [4] were designed to improve the speed of sequence alignment, and are capable of producing optimal and near-optimal alignments in a fraction of the time when compared to Smith-Waterman. However, the improvements in running time for BLAST have not stood up to the accelerated growth of experimental (query) and known (reference) sequence data driven by next-generation sequencing [5]. Genome and metagenome annotation pipelines are therefore left with a few options: (*i*) search against smaller reference databases, (*ii*) use faster, but less sensitive, algorithms to assess sequence similarities [6–8], or (*iii*) invest in computing hardware specifically designed to improve BLAST searches such as GPGPU systems [9] and/or large CPU-rich clusters.

We present a method for hierarchical cluster homology searching, which improves the speed of amino acid sequence homology searches with a minimal decrease in sensitivity and specificity. In general terms, a hierarchical cluster homology search will first search query sequences against a clustered database (e.g. UniRef50 [10]) to identify: (*i*) the query sequences with a match to a cluster representative sequence and (*ii*) the subject sequences belonging to all clusters hit by query sequences. A second homology search is then performed between query sequences with a hit in the first search against subject sequences belonging to the clusters with a hit in the first search. Searching against a subset of the subject sequences in the original (pre-clustered) database results in a linear decrease in search time by passing over subject sequences that would likely not produce a significant alignment. Importantly, this strategy results in sequence alignments and alignment statistics – including E values (expectation values) – that are nearly identical to a BLASTp homology search against the entirety of the original database.

## Implementation

RUBBLE (Restricted clUster BLAST-Based pipeLinE) is a hierarchical cluster protein-protein BLAST (BLASTp) pipeline written in Perl that wraps NCBI BLASTp and is available on GitHub (https://github.com/dnasko/rubble). A typical protein homology search with RUBBLE requires running only one script (`rubble.pl`), which was designed to be very similar to running a command-line BLASTp. Unlike BLASTp, RUBBLE requires that the user provide not only a reference database, but also a clustered version of that database (Fig. 1, part A).

**Figure 1:**
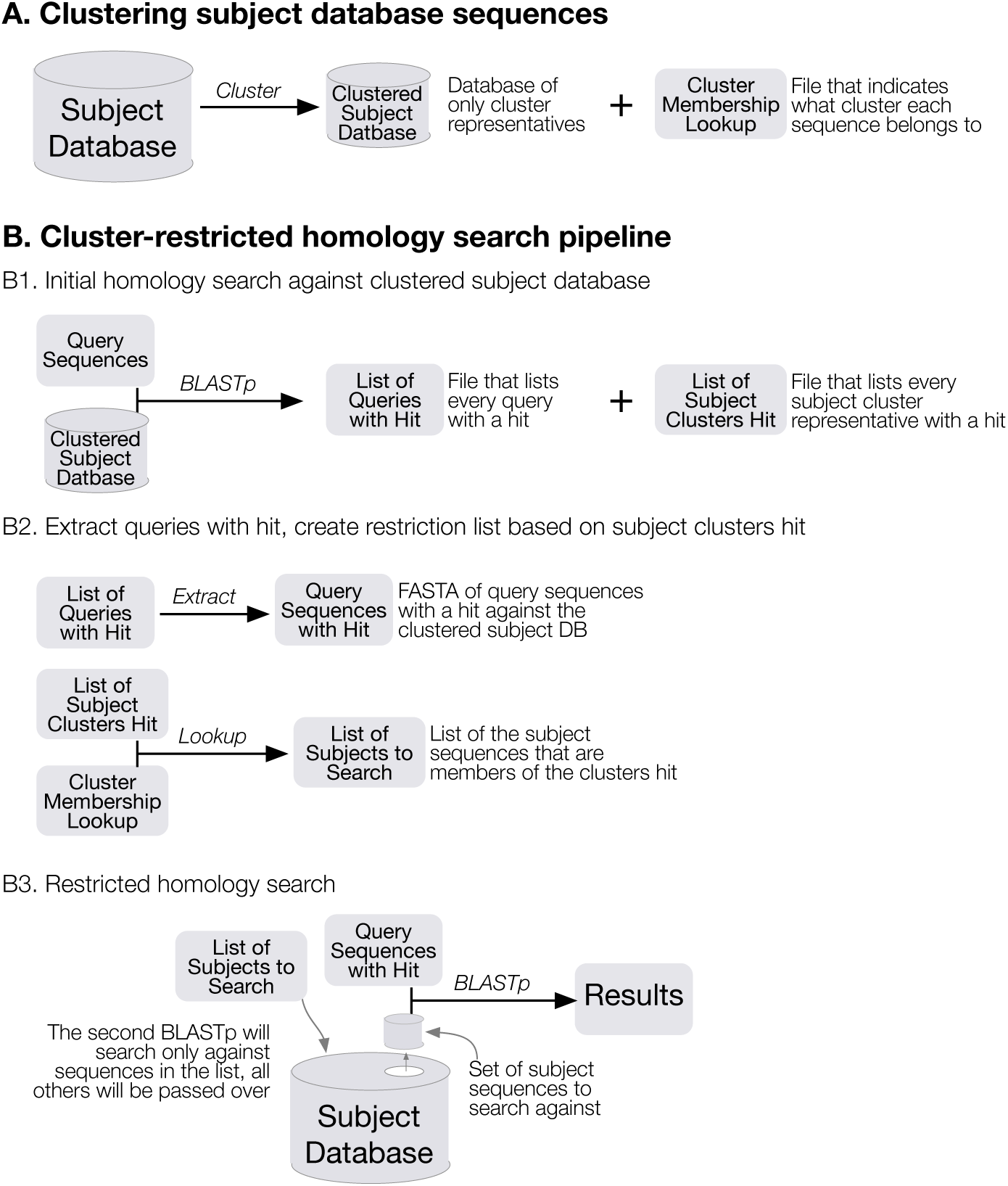
The hierarchical cluster BLASTp workflow, which yields near identical results from a BLASTp search.

Briefly, a RUBBLE homology search will (Fig. 1, part B): (*i*) use BLASTp-fast (`blastp with option`-`task blastp-fast`, a feature available since BLAST+ 2.2.30) [11] to search a set of queries against the database of cluster representatives, (*ii)* extract query sequences that produced an HSP (High-scoring Segment Pair), (*iii*) create a list of subject sequences contained in all of the clusters that had a representative sequence produce an HSP, and finally (*iv*) perform a BLASTp search of the query sequences with a match in the first search against the subject sequences belonging to clusters with a hit from the first search. The second search uses the pre-clustered database (i.e. the original database) as an input, but will be restricted to search against only the subject sequences belonging to clusters with a hit from the first search (using the -`seqidlist` parameter in BLAST). A more detailed explanation is presented below.

### RUBBLE Database Construction

A RUBBLE reference database can be made from any collection of proteins. Given an arbitrary protein database (e.g. Swiss-Prot [12]) users must first cluster this database at a low identity threshold (e.g. 50% or 60%). Next a cluster membership lookup file must be created containing two columns: the sequence ID of the cluster’s representative and the sequence’s ID. This lookup file is crucial as it is used to create the list of the subject sequences to be searched against in the second BLASTp. For example, if subject sequence ‘A’ is hit in the first search and subject sequence ‘A’ is the representative sequence for sequences ‘B’, ‘C’, and ‘D’, then the lookup file will indicate that sequences ‘A’, ‘B’, ‘C’, and ‘D’ will be included in the list of restricted subject sequences for the second search against the pre-clustered database.

Two scripts have been written to build RUBBLE databases from either a custom database (`build_custom_rubble_database. sh` – requires CD-HIT [13]) or from UniRef100 [10] (`build_uniref_rubble_databases.pl`). Building a RUBBLE database from UniRef100 is especially easy because UniProt releases an additional instance of UniRef that is clustered at 50% identity (UniRef50). This saves a great deal of time as it bypasses the need to cluster the database.

Both the clustered and pre-clustered reference BLAST databases are then built using `makeblastdb`. In order to permit the passing of a restriction list to the second search the pre-clustered database needs to be built with the -`parse_seqids` option. Doing so will cause `makeblastdb` to create not only the usual ‘.phr’, ‘.pin’, and ‘.psq’ files, but also ‘.pog’, ‘.psd’, and ‘.psi’ files, which increases the cumulative file size of the database.

### Protein Homology Search with RUBBLE

The RUBBLE protein homology search pipeline can be broadly divided into three steps: (*i*) the initial homology search against a clustered subject database (Fig. 1, part B1); (*ii*) the extraction of query sequences with a hit in the first search and the creation of a list of subject sequences to search against for the second homology search (Fig.1, part B2); and finally (*iii*) the second (restricted) homology search of queries with a match in the first search against subject belonging to clusters with a match in the first search (Fig. 1, part B3).

RUBBLE performs an initial homology search of all query sequences against the clustered reference database (composed of cluster representative sequences) using BLASTp-fast (`blastp` with option –`task blastp–fast`, a feature available since BLAST+ 2.2.30) [11]. This initial search is very fast, as the clustered database should be much smaller than the pre-clustered database (e.g. UniRef50 is 20% the size of UniRef100 as of April 2017). The initial search yields High-scoring Segment Pairs (HSPs) and generates a list of query sequences with a match and a list of cluster representative sequences that were hit by a query.

RUBBLE will then create two files for the second, restricted, search. A restricted query file is created by extracting the header information and sequences of the query sequences with a hit in the first homology search from the original query FASTA file. RUBBLE also identifies the subset of sequences from the pre-clustered database to be searched against in the second homology search. Every subject sequence from the initial homology search that is included in an HSP is a cluster representative sequence. RUBBLE will take the list of subject sequences with a hit from the BLASTp-fast search, use the cluster membership lookup file, and expand this list to include all member sequences from each cluster that had a hit. Specifically, assume subject sequence ‘A’ is hit and subject sequence ‘E’ is not hit in the first search against the clustered database. The cluster membership lookup file indicates that subject sequence ‘A’ is the representative sequence for the cluster containing subject sequences ‘B’, ‘C’, and ‘D’ and that subject sequence ‘E’ is the representative sequence for the cluster containing subject sequence ‘E’, ‘F’, and ‘G’. Because ‘A’ was hit in the first search, but ‘E’ was not, only ‘A’, ‘B’, ‘C’, and ‘D’ will be included in the restricted subject sequences list for the second search against the pre-clustered database (Fig. 1, part B3).

RUBBLE then performs a BLASTp homology search of the restricted query list against the pre-clustered reference database with the restricted subject sequences list (using option -`seqidlist`). By using this restriction list the RUBBLE pipeline is able to reduce the number of subject sequences searched against in the database by 80% to 95%.

## Results and Discussion

The research and development of RUBBLE was the result of a necessity to reduce the computational demand for protein homology searches without compromising the accuracy of results. As the current (and foreseeable [14]) gold standard of accurate and fast protein homology searches is BLASTp, RUBBLE and additional BLAST-like alternatives were tested and compared against the results of BLASTp.

Shotgun metagenomics has perhaps had the greatest impact on investigations of viruses in the environment. There are an estimated 10^31^ free viruses globally [15] making them the most abundant biological entity on the planet, often exceeding bacterial abundance by an order of magnitude [16]. Characterizing viral communities using a single marker gene (akin to rRNA genes in prokaryotes and eukaryotes) is not possible as viruses are polyphyletic [17] and encode modular genomes [18]. Thus, there is a strong desire for shotgun metagenomics in viral ecology as it can randomly sequence all parts of viral genomes from environmental samples. Viral metagenomes are particularly hard to analyze because of the need for distant homology matches and incomplete databases [19]. Thus, evaluating shotgun viral metagenomes serves as a “worst case” for measuring the accuracy of protein alignment programs.

Two assembled shotgun viral metagenomes were used as query datasets in the analysis: an aquatic virome from water collected near the Smithsonian Environmental Research Center (SERC); and a soil virome from Kellogg Biological Stations (KBS). Both of these viromes were searched against three subject databases: UniRef100 [10], Metagenomes Online (MgOl) [19], and Swiss-Prot [12]. More details on the query datasets and subject databases are provided in the data and materials section.

### Comparing RUBBLE with BLASTp-fast

RUBBLE was evaluated against BLASTp-fast, an alternative BLASTp mode that uses longer words for faster seed matching. BLASTp-fast is available for use in all versions of BLAST+ greater than 2.2.30 and allows for faster BLASTp searches. Among current BLAST-like homology search tools, RUBBLE and BLASTp-fast are the most similar to standard BLAST search because they are capable of producing all of the HSP statistics associated with a normal BLAST search (e.g. bit score and E value). Because of this, a stricter set of criteria may be used when comparing RUBBLE and BLASTp-fast with standard BLASTp. Thus, a true positive match between a RUBBLE or BLASTp-fast HSP and a standard BLASTp HSP required a “strict” match in that all of the information match exactly, i.e. all twelve fields in a standard tabular BLAST output (–`outfmt 6`) must match exactly, specifically: `qseqid, sseqid, pident, length, mismatch, gapopen, qstart, qend, sstart, send, evalue, bitscore`.

RUBBLE was able to achieve higher sensitivity (recall: the ability to find all true positives) than BLASTp-fast in each of the six trials performed (Additional File 1). The recall rate for RUBBLE remained stable even when fewer query sequences had a hit to the database, while the recall rate for BLASTp-fast consistently fell when fewer query sequences had hits (e.g. only 10% of SERC ORFs match to Swiss-Prot, RUBBLE recall = 0.95; BLASTp-fast recall = 0.61). The specificity rates (precision: the ability to find true negatives) were consistently high for both RUBBLE and BLASTp-fast, implying neither tool reported many HSPs not identified by BLASTp.

To better understand the performance of RUBBLE and BLASTp-fast based on HSPs identified at various levels of significance, recall rates for each query’s 50 most significant BLASTp HSPs were calculated for the SERC dataset against the three tested reference databases (UniRef, Swiss-Prot, and MgOl) (Fig. 2). Among the top 50 BLASTp HSPs, RUBBLE attained consistently high recall rates (mean recall = 0.97). RUBBLE actually achieved higher recall rates as HSP rankings increased (became less significant). In contrast BLASTp-fast had mean recall rates of 0.77 with lower recall rates as HSP ranks increased.

**Figure 2:**
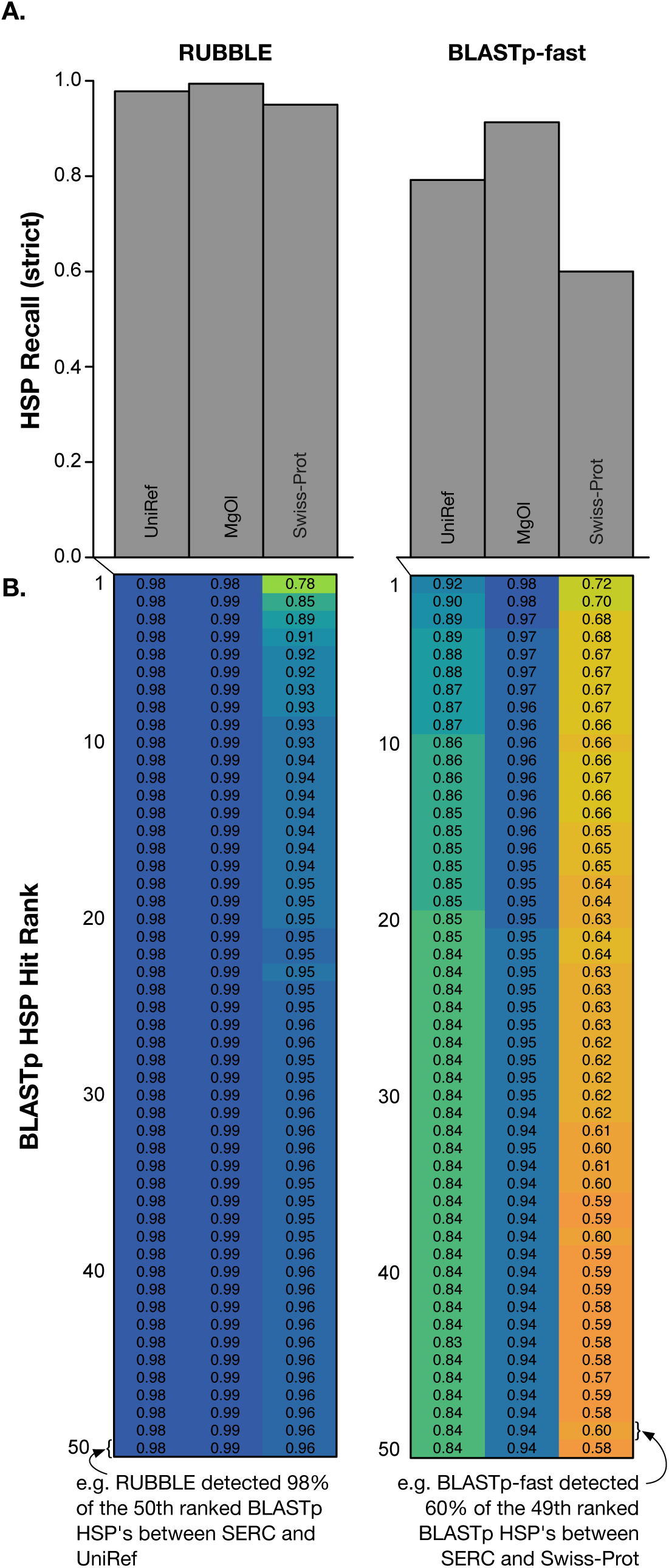
RUBBLE produced more alignments that were identical to BLASTp results than BLASTp-fast. SERC ORFs were searched against three separate databases (UniRef, MgOl, and Swiss-Prot) using RUBBLE and BLASTp-fast. (A) Histogram of recall rates (sensitivity) measured when comparing HSPs produced by RUBBLE and BLASTp-fast with BLASTp (gold standard). A true positive match between a RUBBLE or BLASTp-fast HSP with a BLASTp HSP required that all information contained in the HSPs matched exactly with a BLASTp HSP (e.g. e-value, bit score, coordinates, etc.), this is considered a “strict” match. RUBBLE achieved a higher recall rate than BLASTp-fast against all databases. (B) The recall rates of each of the top-50 BLASTp HSPs for RUBBLE and BLASTp-fast. Each row is the HSP rank from a BLASTp search and the value in each box indicates how often an HSP from that tool against that database matched a BLASTp HSP of that rank. Box color corresponds to higher fractions (dark blue, better) and lower fractions (orange, worse).

### Comparing RUBBLE with BLASTp Alternatives

BLASTp alternatives, such as those tested here (BLAT, DIAMOND, LAST), are able to produce statistics similar to those output by BLASTp such as E value, and bit score [8, 20, 21]. However, the ways in which these values are calculated can differ greatly from BLASTp, making it difficult and impractical to compare these results. For this evaluation, the HSPs produced by RUBBLE and the other BLASTp alternatives need only fit a “relaxed” match, i.e. they need only find all of the query-subject pairings that BLASTp found, regardless of the alignment statistics. For example, if BLASTp found Query A hit Subject B at 100% identity, and LAST found Query A hit Subject B at 90% identity, then this would still be considered a valid “relaxed” match despite the discrepancy between percent identities.

With relaxed matching criteria, RUBBLE still outperformed BLASTp-fast in terms of recall and outperformed the BLAT, DIAMOND and LAST by even wider margins (Table 1). In every trial, RUBBLE was able to produce a set of HSP results more similar to BLASTp than other tools tested (Fig. 3) despite differences in subject database size and number of query sequences with a hit.

**Table 1:** A relaxed comparison of RUBBLE and BLASTp-like programs

| Program | Mean Recall | Mean Precision | Time reduction* |
| --- | --- | --- | --- |
| RUBBLE | 0.97 | 0.99 | 12 |
| BLASTp-fast | 0.78 | 0.96 | 12 |
| BLAT | 0.19 | 0.33 | 178 |
| DIAMOND | 0.07 | 0.98 | 1,995 |
| LAST | 0.07 | 0.68 | 34,858 |

\* Time reduction = Mean program CPU time / BLASTp CPU time

**Figure 3:**
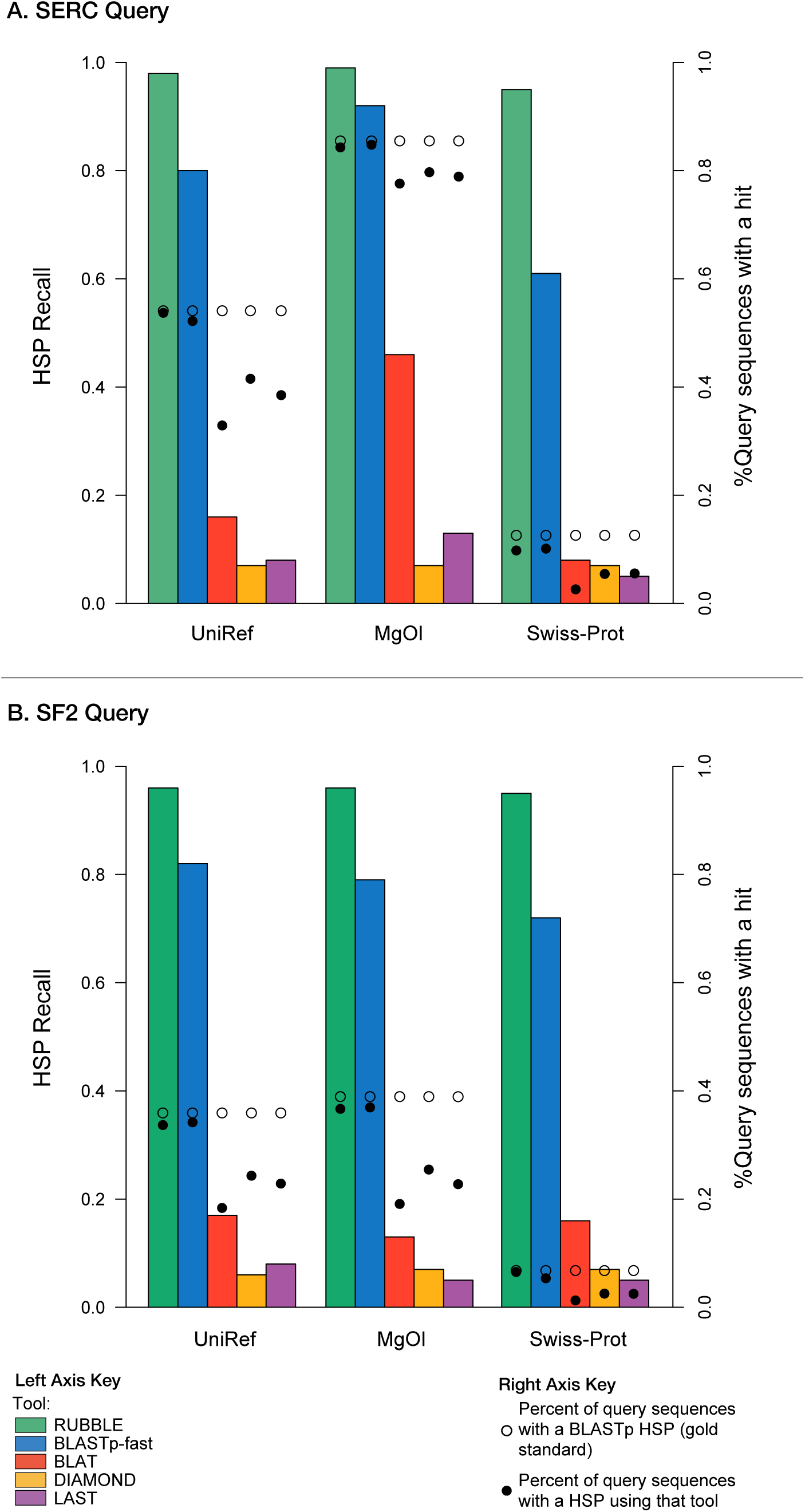
RUBBLE is better at producing HSPs identical to BLASTp than alternative homology search tools. The SERC (A) and SF2 (B) datasets were searched against UniRef, MgOl, and Swiss-Prot using RUBBLE, BLASTp-fast, BLAT, DIAMOND and LAST. The clustered bars indicate the recall rate of each tool’s results relative to BLASTp. A true positive match between a RUBBLE, BLASTp-fast, BLAT, DIAMOND or LAST HSP with a BLASTp HSP required only that a given the query-subject pairing matched a BLASTp query-subject pairing (i.e. the E value, bit score, percent identity, etc. did not have to match). Again, RUBBLE outperformed BLASTp-fast and outperformed the three BLASTp-alternative tools by an even wider margin. The black points correspond to the right-side Y-axis. The filled points indicate the percent of query sequences with a hit to each database using that tool, while the empty points indicate the percent of query sequences with a hit to each database using BLASTp (gold standard). HSPs from RUBBLE strongly matched those from BLASTp when many queries have a hit to a database (e.g. SERC against MgOl) and when few queries have a hit to a database (e.g. SF2 against Swiss-Prot).

Additionally, the HSPs that were missed by RUBBLE were often of lesser significance in terms of E value than the other programs tested (i.e., RUBBLE rarely missed HSPs that were more significant) (Fig. 4). As E value is calculated based on database size it is only possible to estimate this value, by using a large database (e.g. UniRef100), thus the estimates shown in Fig. 4 are all fairly conservative (i.e. likely higher/less significant than one would expect against smaller databases).

**Figure 4:**
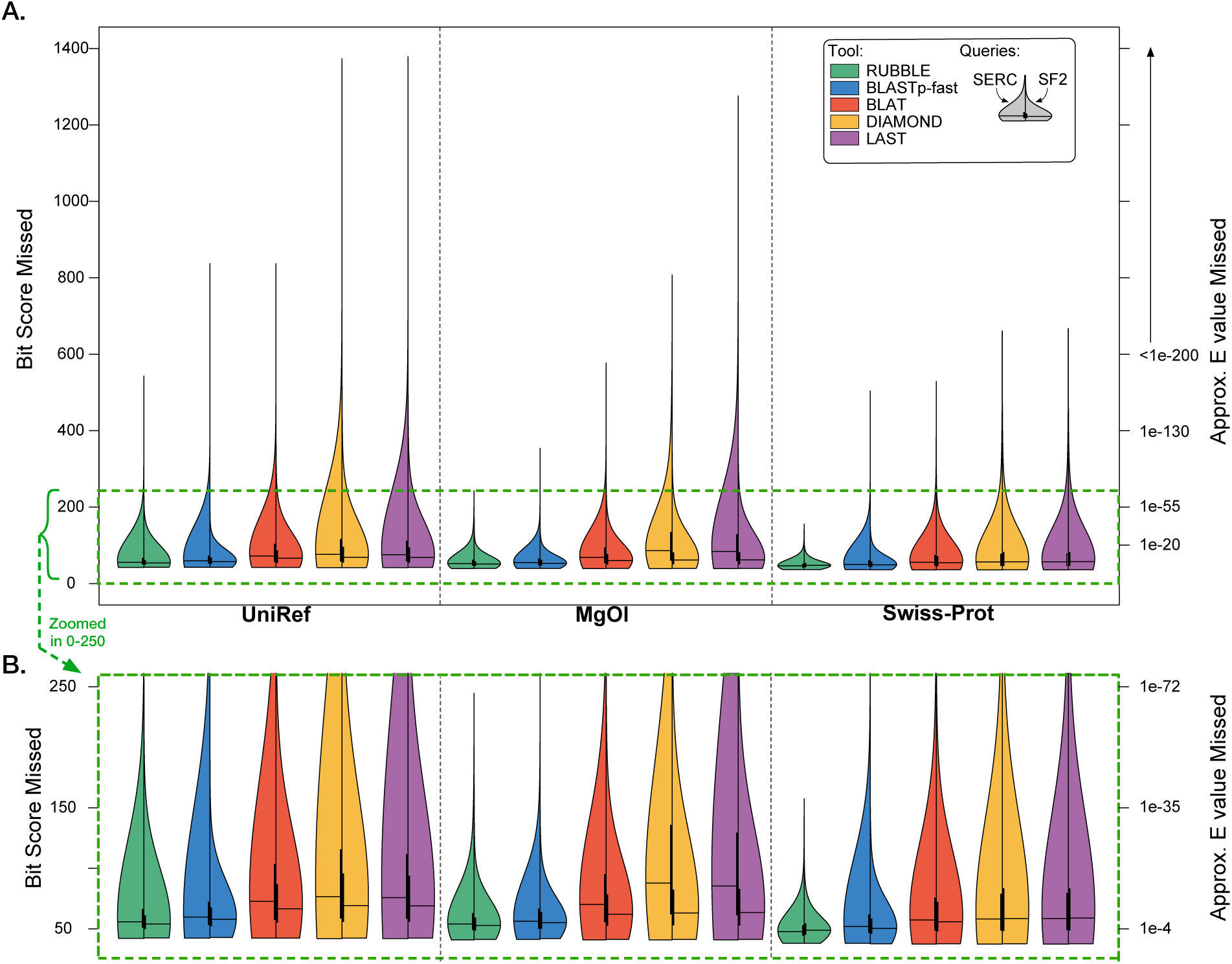
Bit scores of BLASTp HSPs that RUBBLE and BLASTp-fast missed were smaller than bit scores missed by BLAT, DIAMOND and LAST; i.e. RUBBLE and BLASTp-fast are more likely to miss less significant HSPs than BLAT, DIAMOND or LAST. (A) Split violin plots of bit scores of BLASTp HSPs missed by each algorithm searching SERC (left split) and SF2 (right split) against each database. The top (A) shows the whole range of bit scores, the bottom (B) is the same plot zoomed to show bit scores missed from 0-250. An approximate E value for each bit score is provided on the right-side Y-axis. As E values depend on database size, these values are conservative estimates based on the largest database (UniRef).

All of the BLASTp-like tools required less CPU time compared BLASTp (“time reduction” field in Table 1). Both RUBBLE and BLASTp-fast ran approximately 12 times faster than BLASTp. For RUBBLE the speed-up ranged from 6 to 20 times faster than BLASTp (Additional Files 1 and 2). BLAT ran approximately 170 times faster than BLASTp (range = 141-250), DIAMOND ran approximately 2,000 times faster (range = 1,400-3,000), and LAST ran approximately 35,000 times faster (range = 6,000-68,000). No significant correlations were detected between time reduction and any variable, however some trends did emerge. For RUBBLE the time reduction appeared to correlate negatively with the percentage of query sequences producing a HSP (i.e. more queries with HSPs likely leads to a smaller time reduction; fewer queries with HSPs likely leads to a larger time reduction). For BLAT, DIAMOND and LAST the speed-up appeared to correlate positively with the size of the reference database size. This is because the algorithms that these programs are based on have a non-linear dependence on database size, unlike BLASTp.

## Conclusion

We have presented an implementation of cluster-restricted BLASTp called RUBBLE. This novel method allows for protein homology searches that are 10 to 20 times faster than BLASTp. Through validation with two viromes and three subject databases, of varying sizes and compositions, we have demonstrated that RUBBLE consistently produces results that are nearly identical to BLASTp. Additionally, RUBBLE outperformed currently available BLASTp alternatives BLASTp-fast, BLAT, DIAMOND and LAST in terms of recall and precision. While the 10X to 20X reduction in CPU time is modest in comparison to BLAT, DIAMOND and LAST (capable of achieving reductions of >10,000X), RUBBLE provides a method to reduce CPU demand while producing protein homology results with high fidelity, not achievable with other BLAST-like alternatives.

RUBBLE will be maintained and kept up-to-date on GitHub as future versions of commandline NCBI BLAST update. Additionally, we may test the cluster-restricted BLAST method with nucleotide-protein BLAST (BLASTx) and nucleotide-nucleotide BLAST (BLASTn).

## List of Abbreviations

BLAST: basic local alignment search tool
BLAT: blast-list alignment tool
ESS: environmental shotgun sequencing
HSP: high-scoring segment pair
RUBBLE: restricted cluster blast-based pipeline
SERC: Smithsonian Environmental Research Center

## Declarations

### Ethics approval and consent to participate

Not applicable.

### Consent for publication

Not applicable.

### Availability of data and material

The query sequences used in this analysis were predicted peptide ORFs (open reading frames) from viral shotgun metagenomes (viromes). The SERC virome sampled aquatic viruses from the Chesapeake Bay and was sequenced using Illumina Hi-Seq producing ca. 77 million paired reads and are available on the NCBI Sequence Read Archive (https://www.ncbi.nlm.nih.gov/sra) accession number: SRR4293227. As this analysis required many BLASTp searches against large databases it was decided that a smaller subset of the whole SERC dataset would be used for analysis. A random 10% of reads were pulled from the whole data set using a Perl parser (https://github.com/dnasko/rubble/blob/master/manuscript/sampler.pl). These reads were merged using FLASh [22] ver. 1.2.6 and assembled with SPAdes [23] ver. 3.6.2 (using “only assembler”). ORF’s were predicted from each contig using Metagene Annotator [24].

The second query dataset used in this analysis, SF2, was a soil virome collected from free viruses at the Kellogg Biological Station (University of Michigan, Hickory Corners, Michigan) and sequenced using 454 FLX Titanium, producing ca. 1 million reads. These reads were filtered for artificial duplicates using CD-HIT-454 [25] and assembled using SPAdes (using “only assembler”). Again, ORF’s were predicted using Metagene Annotator.

### Authors’ contributions

D.J.N and S.W.P. designed research; D.J.N. performed the research; D.J.N. wrote the software; D.J.N. and S.W.P. wrote the paper; D.J.N., B.D.F., S.W.P., and K.E.W. revised the paper.

## Acknowledgements

Support from the University of Delaware Center for Bioinformatics and Computational Biology Core Facility and use of the BIOMIX compute cluster was made possible through funding from Delaware INBRE (NIGMS P20 GM103446) and the Delaware Biotechnology Institute.

Additionally, this research was supported in part through the use of Information Technologies (IT) resources at the University of Delaware, specifically the high-performance computing resources.

## Additional Files

**Additional File 1:** Strict comparison of RUBBLE and BLASTp-fast.

**Additional File 2:** Relaxed comparison of RUBBLE, BLASTp-fast and BLAST-like programs.

